# Microglia motility depends on neuronal activity and promotes structural plasticity in the hippocampus

**DOI:** 10.1101/515759

**Authors:** Felix C. Nebeling, Stefanie Poll, Lena C. Schmid, Manuel Mittag, Julia Steffen, Kevin Keppler, Martin Fuhrmann

## Abstract

Microglia, the resident immune cells of the brain, play a complex role in health and disease. They actively survey the brain parenchyma by physically interacting with other cells and structurally shaping the brain. Yet, the mechanisms underlying microglia motility and their significance for synapse stability, especially during adulthood, remain widely unresolved. Here we investigated the impact of neuronal activity on microglia motility and its implication for synapse formation and survival. We used repetitive two-photon *in vivo* imaging in the hippocampus of awake mice to simultaneously study microglia motility and their interaction with synapses. We found that microglia process motility depended on neuronal activity. Simultaneously, more dendritic spines emerged in awake compared to anesthetized mice. Interestingly, microglia contact rates with individual dendritic spines were associated with their stability. These results suggest that microglia are not only sensing neuronal activity, but participate in synaptic rewiring of the hippocampus during adulthood, which has profound relevance for learning and memory processes.

## Introduction

Microglia are cells of the innate immune system in the central nervous system (CNS). Under physiological conditions microglia are sessile, highly ramified and plastic cells that constantly scan their environment with fine processes, referred to as microglia motility^1, 2^. This remarkable degree of cellular plasticity enables microglia to detect and quickly react to even subtle changes of tissue homeostasis within their microenvironment - a process that has been shown to be ATP-dependent^2^. While microglia have been linked to a number of neurodegenerative diseases such as Alzheimer’s disease^3, 4^, still little is known about their physiological function under healthy conditions^5, 6^. Already during the discovery of microglia motility the hypothesis was tested whether their process motility related to neuronal activity. Interfering with neuronal activity *in vivo* using Tetrodotoxin (TTX) and Bicuculine led to inconsistent finding^1^. It might be, that these results were confounded by the usage of anesthetics like isoflurane, which was shown to suppress microglia baseline motility *in vitro^7^*. Whether this holds true for the *in vivo* situation remains an open question. Evidence from Ca^2+^-imaging in zebrafish suggests a reciprocal relationship of microglia motility and neuronal activity showing decreased Ca^2+^-transient frequencies upon microglia contact of neurons^8^. Further evidence for a dependency of microglia process motility and neuronal activity derived from studies investigating the relationship of dendritic spine formation and elimination using two-photon *in vivo* imaging. Lowering neuronal activity by applying hypothermia and TTX reduced the frequency of microglia contacts with dendritic spines in the visual and somatosensory cortex. However microglia motility was not directly measured in this study^9^. Moreover, in the visual cortex microglia motility was reduced in mice that were kept in darkness, whereas it was increased in mice re-exposed to light after a dark period^10^. The same study found that microglia contact rates were increased before dendritic spines were eliminated during a critical period of developmental plasticity, suggesting a role for microglia in shaping post-synapses. Also during early development in the mouse cortex, microglia contacts at dendrites induced the formation of new dendritic filopodial spines, yet only in awake, not anesthetized mice^11^. Additionally removal of microglia from the brain in adult mice affected the turnover rate of dendritic spines in the motor cortex and motor learning induced formation of new spines was reduced, in an BDNF dependent manner^12^. The main brain region involved in learning and memory processing is the hippocampus. Whether microglia in the hippocampus have similar functions in dendritic spine formation and elimination as in the cortex remains unresolved. Yet, recent data from *in vitro* studies showed a relationship of long-term potentiation and spine contact rates by microglia in the hippocampus^13^. Moreover, pre-synapses but not post-synapses were eliminated by trogocytosis (non-apoptotic uptake of membrane components by immune cells), whereas filopodial spines seemed to be induced by microglia during development in hippocampal slice cultures^14^. Microglia motility has not been examined *in vivo* in the hippocampus up to now and recent findings implicate a more heterogeneous role of microglia throughout different brain regions then previously assumed^15^. It is therefore important to decipher whether microglia motility in the hippocampus is different from the cortex. Furthermore, the interplay between microglia and synapses is of particular interest as microglia are by far the most plastic cells in the CNS and by engaging in contact with synapses they could have the potential to enhance and weaken synaptic transmission as well as affect structural plasticity of spines, both during health and disease. Whereas microglia have been shown to be critical for synaptogenesis and synaptic integrity during developmental stages^10, 11, 16, 17^, only little is known about their contribution to structural plasticity in the hippocampus during adulthood. Thus, it is critical to understand the nature and functional properties of microglia synapse interactions, especially during adulthood, as the role of microglia in CNS homeostasis might differ between CNS development and adulthood^18^. For that reason we set out to investigate microglia motility under healthy conditions and in the absence of anesthesia in the hippocampus of awake, adult mice while examining to what extend baseline surveillance of microglia is dependent on neuronal activity. Additionally, we address the question whether microglia play a role in dendritic spine formation and elimination in the hippocampus during adulthood supporting the lifelong rewiring of the hippocampal neuronal micro-network.

## Results

### Increased microglia motility in awake mice

To address the relationship of neuronal activity and microglia motility in the hippocampus *in vivo*, we used double transgenic mice expressing green fluorescent protein (GFP) in microglia^19^ and yellow fluorescent protein (YFP) in neurons^20^. Mice were heterozygous for both transgenes Thy1-YFP^+/-^::CX3CR1-GFP^+/-^ and between 7-10 months of age. To access the hippocampus we unilaterally removed parts of the somatosensory cortex and implanted a metal tube, sealed with a glass cover-slip at the bottom^21^. After 3-4 weeks of recovery, two-photon *in vivo* imaging started (Fig. 1a,b). Microglia motility was investigated in anesthesia, in awake mice running on a circular treadmill and in awake mice with topical TTX application (Fig. 1a). Microglia motility was measured by acquiring z-stacks spanning 100 μm in *stratum radiatum* of the dorsal hippocampus (Fig. 1b). Microglia process turnover rate was calculated as previously described^22^. The lost and gained proportion of microglia processes from one to the next imaging time point was calculated in a two-channel color overlay image (Fig. 1c; Material and Methods). Under isoflurane anesthesia (1.0% in O_2_), microglia showed reduced basal motility compared to microglia in awake mice (Fig. 1d; 60.2% ± 2.7% anesthesia vs. 69.9% ± 2.8% awake; Supplementary Videos S1+S2). Isoflurane mainly acts by prolonging GABAergic transmission thus inhibiting neuronal activity^23–25^ (additionally see Supplementary Fig. 1). Therefore, we investigated to what extend the increased level of microglia motility in awake mice was due to an increase of neuronal activity. To address this question, we topically applied TTX (50 μM), a highly potent blocker of voltage-gated sodium channels, to inhibit action potential firing in the hippocampus. This pharmacological manipulation significantly decreased microglia motility in awake mice, even below the levels of mice in anesthesia (Fig. 1d; 69.9% ± 2.8% awake vs. 52,8 ± 1.2% awake plus TTX). These results suggest a relationship of microglia motility with neuronal activity in the hippocampus of awake mice.

**Figure 1.**
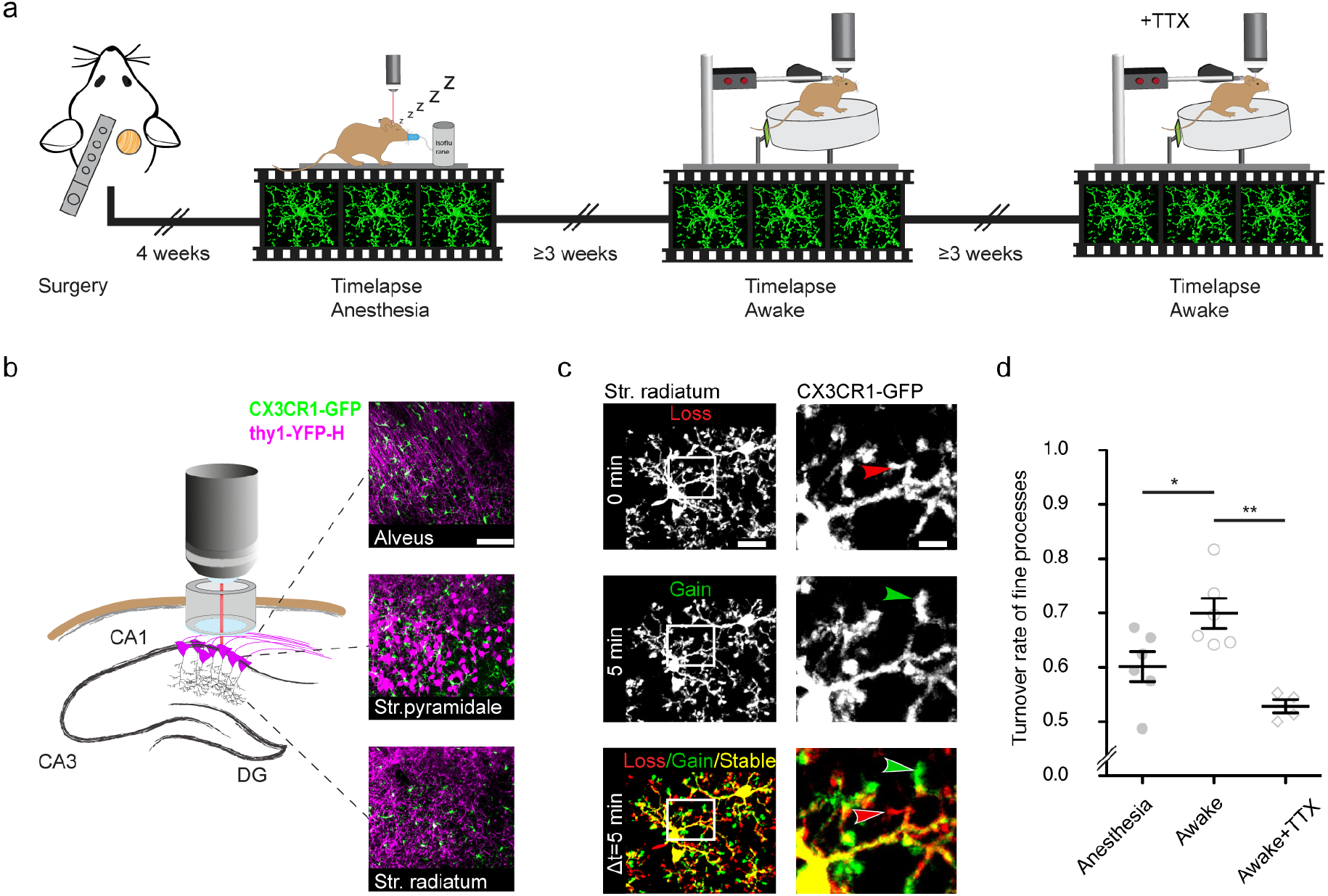
Increased microglia motility in the hippocampus of awake mice. **a** Experimental paradigm of repetitive microglia imaging in the same mice under varying conditions (anesthesia, awake, awake+TTX). **b** Illustration of the accessible hippocampal layers in CA1 through a chronic hippocampal window. CX3CR1-GFP::thy-1-YFP-H transgenic mice allowed for simultaneous visualization of microglia (green) and neurons (magenta). **c** Representative image of a microglia cell and its processes in an awake mouse. Subsequent time points (0 min, 5 min) were overlaid (Δt = 5 min) to measure gained (green arrow), lost (red arrow) and stable (yellow area) processes. **d** Turnover rate (TOR) of microglia fine processes (i.e. motility) at varying conditions. n=6 mice anesthesia, awake; n=4 awake+TTX; one-wayANOVA with Bonferroni’s multiple comparison test, F(2,13) = 10.22; adjusted p = 0.0291 (anesthesia vs. awake), p = 0.0014 (awake vs. awake + TTX). Scale bars: (b) 100μm, (c) 10μm, 2μm; * p < 0.05, ** p < 0.01.

### Microglia motility in awake mice depends on neuronal activity

Under physiological conditions microglia only express a very limited number of voltage-gated sodium channels^26^. Nevertheless, a direct effect of TTX on microglia cannot be excluded entirely. To investigate this and to further clarify whether microglia motility in the hippocampus is depended on neuronal activity, we specifically interfered with neuronal activity. Therefore, we chose a chemogenetic approach distinctly silencing hippocampal neurons *in vivo*. To this end, we injected two adeno-associated viruses (AAVs), one expressing Cre-recombinase under the neuron-specific Ca^2+^/calmodulin-dependent protein kinase II (CaMKII) promoter (AAV9-CaMKII-Cre, Addgene) and another loxP-flanked version of an inhibitory G-protein coupled receptor (hM4D(Gi)) under the neuronal synapsin I promoter tagged with the fluorescent reporter mCherry (AAV2-hsyn-DIO-hM4D(Gi)-mCherry^27^). The inhibitory G-protein coupled receptor hM4D(Gi) is a DREADD (designer receptor exclusively activated by designer drug) that upon binding of the drug clozapine N-oxide (CNO), leads to silencing of transfected neurons^28^. To simultaneously silence CA1 pyramidal neurons and their major input coming from CA3 neurons via the Schaffer collaterals, we injected into both regions, CA1 and CA3 (Fig. 2a). The expression of the inhibitory DREADD was confirmed by post-hoc immunohistological stainings of transfected hippocampi (Supplementary figure S2). We analyzed microglia motility in the same awake mice with (CNO) and without (vehicle) the specific silencing of neurons in CA1 (Fig. 2b). The level of microglia motility in mice treated with vehicle was comparable to the first set of mice recorded under awake conditions (Fig. 1d vs. Fig. 2c; 69.9% ±2.8% vs. 71.2 ±1.5%), reproducing the high level of microglia motility in the hippocampus of awake mice. However, a treatment with CNO resulted in a significant reduction of microglia motility, 30-90 min after application (Fig. 2c; 71,2 ±1.5% vehicle vs. 58,7 ±1,8% CNO). These results indicate that microglia motility depends on neuronal activity in the hippocampus of awake mice.

**Figure 2.**
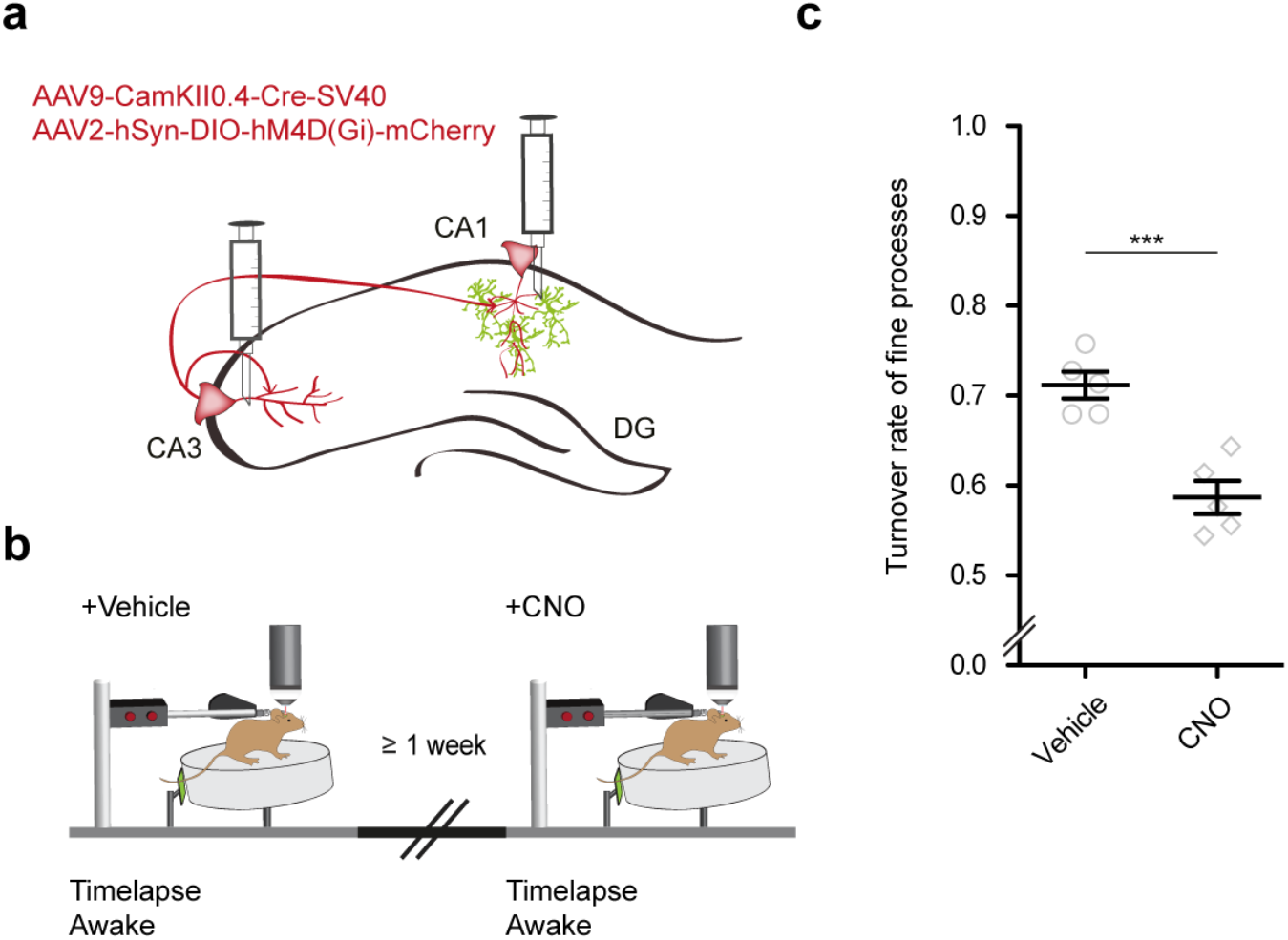
Microglia motility in awake mice depends on neuronal activity. **a** Injection of two AVVs into the CA1 and CA3 region of the hippocampus with CamKII-driven Cre-recombinase expression in neurons (1^st^ AAV). This allows for the recombination of loxP-flanked hM4D(Gi) and expression of the DREADD under human synapsin1 promoter (hSyn) in these neurons (2^nd^ AAV). **b** The same mice were imaged awake twice, with a time interval of at least one week. Imaging started 40 min. after i.p. injection of either vehicle or CNO. **c** Turnover rate of microglia fine processes in vehicle - and CNO - treated mice; n=5 mice, paired t-test, two tailed, p=0.0048. *** p < 0.001.

### Spine stability is associated with microglia contact rate

Given that the Thy1-YFP^+/-^::CX3CR1-GFP^+/-^ mouse line additionally allows the visualization of neuronal structures, we investigated whether there existed differences in structural plasticity of dendritic spines in anesthetized and awake mice. (Supplementary Fig. 2a). We acquired z-stacks containing radial oblique dendrites in *stratum radiatum* of the hippocampus. Repetitive imaging spaced by two days, first in anesthesia and then awake, we observed a significant increase in spine density as well as significantly higher fractions of gained compared to lost spines in awake mice (Supplementary Fig. 2c-f). These results suggest more rewiring of hippocampal synaptic connectivity in awake mice, underscoring the plasticity of the hippocampus during adulthood. Moreover, a substantial fraction of turnover of dendritic spines occurred spatially clustered along the dendrites of CA1 pyramidal neurons (55% of turnover, see Supplementary Fig.3a-c). Similar findings, have previously been reported for dendrites of excitatory neurons in the somatosensory, motor, visual and retrosplenial cortex^29–31^, underscoring our finding that rewiring of hippocampal neuronal connections happens in close proximity. After identification of changes in the structural plasticity of dendritic spines we investigated, whether microglia were involved in these alterations, as they have previously been linked to both synaptogenesis and synaptic degradation^10–12, 14, 17, 32^. Taking into account our finding that microglia motility depended on neuronal activity, we hypothesized that increased microglia motility would be associated with structural plasticity of dendritic spines in the hippocampus of adult mice. To address this hypothesis, we analyzed the interaction of microglia with individual spines and the dendritic shaft in *stratum radiatum* of the hippocampus *in vivo* (see Supplementary Video 3 for example). Theoretically, after being contacted by microglia, a dendritic spine might either be stable or lost consecutively (Figure 3a). In addition, microglia contact of the dendrite may also lead to the formation of a new spine (Figure 3a). To investigate these possibilities, we carried out a 45-minute time-lapse imaging with 5-minute intervals on the first day and another imaging two days later to associate microglia contact rates with subsequent loss or gain of spines. Indeed, we observed spine loss upon microglia contact (Figure 3b-d) as well as spine formation upon microglia dendrite contact (Figure 3e-g). Quantifying individual contacts of microglia with dendritic spines in anaesthetized and awake mice revealed significantly higher contact rates in awake mice (Fig. 3h; 5.34 ± 0.33h^−1^ in anesthesia vs. 7.33 ± 0.16h^−1^ awake). However, no significant difference between awake and anaesthetized mice was found for contacts of microglia with the dendritic shaft (Fig. 3i; 5.55 ± 0.28h^−1^ in anesthesia vs. 6.22 ± 0.26h^−1^ awake). To find out whether there existed any differences concerning the contact rate of spines between the different categories of stable, lost and gained spines we analyzed their microglia contact rates. Interestingly, the contact frequency of lost spines was significantly increased compared to stable spines (Fig. 3j; stable spines: 5.34 ± 0.33h^−1^ vs. lost spines 8.63 ± 0.81h^−1^). The same was true for gained spines compared to stable spines, indicating that an increased microglia contact frequency is associated with higher structural plasticity of dendritic spines (Fig. 3j; stable 5.34 ± 0.33h^−1^ vs. gained 9.94 ± 0.33h^−1^). Moreover, microglia contacts were more frequent at dendritic shafts on which spines were gained compared to dendritic segments where no spine gain was detectable (Fig. 3j; dendritic shafts without spine gain 5.55 ± 0.28h^−1^ vs. dendritic shafts with spine gain 9.94 ± 0.33h^−1^). Carrying out the same comparisons in awake mice revealed similar changes between groups, yet the overall number of contacts was increased in awake mice (Fig. 3k; stable spines 7.33 ± 0.16h^−1^, lost spines 11.4 ± 0.21h^−1^, gained spines 10.49 ± 0.26h^−1^, dendritic shaft 6.22 ± 0.26h^−1^). Furthermore, microglia contacts of dendritic spines were significantly higher if spine turnover occurred in clusters (Supplementary Fig.3d), suggesting microglia might facilitate spatial reorganization of dendritic spines in the hippocampus.

**Figure 3.**
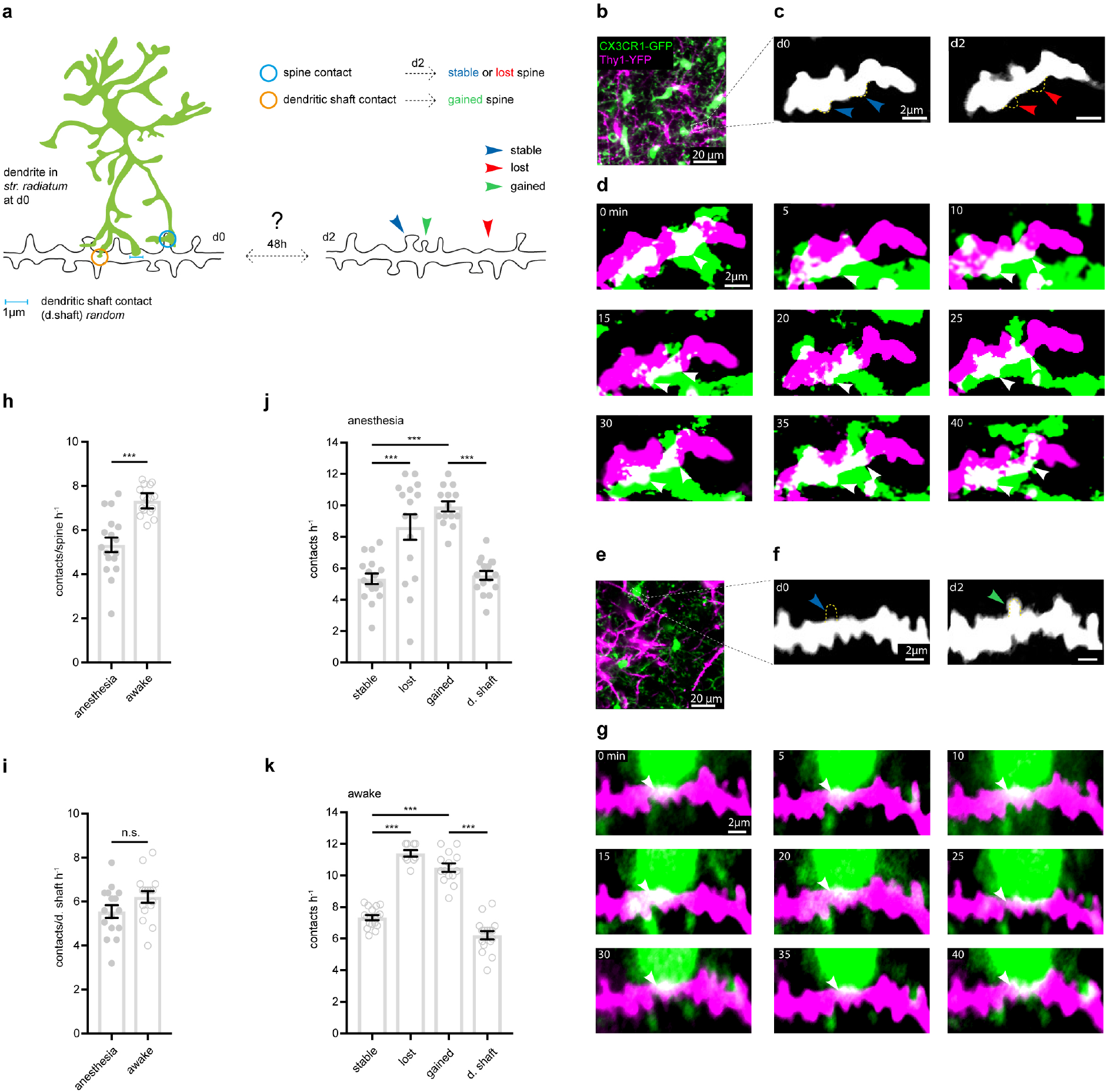
Microglia contact frequency associated with spine stability in the hippocampus. **a** Schematic illustrating the analysis between microglia contacts and spine stability: direct spine contacts (blue circle) can lead to a stable (blue arrow) or lost spine (red arrow) as analyzed two days later, whereas contacts of the dendritic shaft (orange circle) can lead to a gained spine (green arrow) or no change. **b** Image from the hippocampus showing microglia cells (green) and neurons (magenta) in a CX3CR1-GFP::thy-1-YFP-H mouse. **c** Zoom image from (b) illustrating a dendritic segment with two spines (blue arrows, day 0) that are lost on day 2 (red arrows). **d** Time-lapse of microglia processes interacting with the dendrite and spines (white area) in the hippocampus of an awake mouse; arrows (orange) mark individual microglia contacts of spines. **e-g** Analog illustration as (b-d) for a gained spine example (green arrow). **h,i** Direct microglia contacts (h) and contacts along the dendritic shaft (i) in anesthetized versus awake mice unpaired t-test (h,i); p < 0.0001 (h), p = 0.0964 (i)). **j,k** Contact rate in anesthesia (j) and awake (k) at stable, lost and newly gained spines as well as at random locations along the dendrite. One-way ANOVA with Sidak’s multiple comparison test (j,k); F (3, 59) = 20.88 (j) and F (3, 51) = 103.7 (k); Spine contacts averaged over dendrites. with n = 17 stable, 16 loss, 14 gain, 16 d. shaft in anesthesia and n = 16 stable, 9 loss, 14 gain and 16 d. shaft awake in n = 4 mice(h-k); * p < 0.05, ** p < 0.01, ***p < 0.0001.

In summary, these data show that microglia contact rates of spines are increased in awake mice corresponding to a general increase of microglia motility dependent on neuronal activity. Furthermore, the turnover of dendritic spines was associated with a higher microglia contact frequency in the hippocampus of adult mice.

## Discussion

In this work we set out to investigate different functional properties of microglia in the hippocampus of mice *in vivo* by means of two-photon microscopy. We detected elevated motility of microglia fine processes in awake mice compared to mice in anesthesia. We demonstrate a dependence of microglia motility on neuronal activity using pharmacologic and chemogenetic approaches. Regarding the role of microglia in maintaining or destabilizing synapses, we identified an association of microglia contacts of dendritic spines and their stability. Lost spines and dendritic stretches with a newly appearing spine were more frequently contacted by microglia than stable spines. Lowering neuronal activity by anesthesia lowered microglia-spine interaction in general, but did not affect the aforementioned relation. Furthermore, clustered events of spine gain or loss were associated with higher microglia contact rates. These results form the basis for future work unraveling whether microglia-mediated synaptic rewiring is involved in hippocampus-dependent behavior, like spatial navigation, learning and memory.

### Microglia process motility

Recent findings indicate a considerable effect of the volatile narcotic isoflurane on microglia by acting on THIK-1 channels and reducing microglia scanning activity *in vitro^7^*. Here we investigate microglia motility *in vivo*, comparing anesthesia and awake conditions in mice. Microglia fine process motility was reduced in anesthetized mice compared to awake conditions. Our data suggest that the reduction of microglia motility might be due to a decreased level of neuronal activity. Applying TTX to the hippocampus, we observed a markedly reduction of microglia motility in awake mice. However, in previous studies no such effect of TTX on microglia process motility was detectable^1, 9, 33^. Possibly in previous experiments, neuronal activity was already reduced to a certain degree by the usage of anesthetics^1, 9^ or the experiments were performed in slice cultures^33^, where neuronal activity and connectivity is not intact anymore. Another difference between the past *in vivo* studies and the current one is the analyzed brain region. Our study has been carried out in the *stratum radiatum* of the hippocampus, whereas former studies investigated the somatosensory cortex. To rule out a direct modulatory effect of TTX on microglia motility, we used a chemogenetic approach to specifically silence neurons *in vivo*. We found a reduction of microglia motility after silencing neurons in CA3 and CA1 in awake mice to a degree comparable to the changes found between anesthetized and awake animals. Therefore, we conclude a dependency of microglia motility on neuronal activity in the hippocampus of mice. It should however be noted that baseline motility never subsided entirely, but rather was attenuated following pharmacological and chemogenetic manipulations. Although a complete reduction of neuronal activity *in vivo* cannot be achieved, our findings suggest further regulatory mechanisms of microglia motility. In summary, these findings argue for a more cautious use of narcotics in *in vivo* experiments. Especially functional assays of microglia using isoflurane should be interpreted cautiously.

### Microglia contact rates of dendritic spines are increased in awake mice

Up to now, only few studies investigated changes in hippocampal synaptic connectivity *in vivo* by analyzing structural plasticity of dendritic spines. The first study investigated structural plasticity of spines in the hippocampus over hours detecting only minor changes^34^. By investigating structural plasticity over weeks, we could demonstrate synaptic rewiring in adult mice^21^. In addition to excitatory neurons, also some inhibitory neurons exhibit structural plasticity of dendritic spines^35^ supporting the view that synapses in the hippocampus are constantly emerging and eliminated. This capacity might enable the hippocampus to adapt to altered external stimuli and different environments. Indeed, the structural plasticity of dendritic spines in the hippocampus might still be underestimated as recent studies using superresolution approaches revealed high turnover rates of dendritic spines, although the analysis lasted for only days to weeks^36, 37^. However, all previous studies have been carried out in anesthesia. Here we demonstrate structural plasticity of dendritic spines on apical dendrites in CA1 in awake mice, which offers the possibility for future studies to combine this kind of imaging with defined sensory stimuli or behavioral tasks. Indeed, awake mice exhibited a higher spine density after two days compared with anesthetized mice. We interpret the increased spine density after awake imaging most likely as a response to an enriched environment, which has been shown to induce a dendritic spine increase in the hippocampus^38^ and the cortex^39^.

In order to integrate our findings on microglia motility with structural plasticity of dendritic spines, we examined the interaction of microglia processes with dendritic spines. While microglia have traditionally been studied in the context of immune responses and inflammatory processes^40, 41^, a growing amount of evidence shows the importance of microglia for normal CNS maturation, function and homeostasis apart from classical immune reactions^18, 42^. A compelling idea suggests microglia might actively contribute to structural plasticity of pre- and post-synapses. This hypothesis derives from a number of studies showing the necessity of normal microglia function for synaptic integrity and learning dependent synapse formation^10–14, 16, 17, 32, 43–48^. Yet, studies directly monitoring microglia synapse interaction are rather sparse^9–11, 13, 14, 49^. These studies have been performed either in the cortex under anesthesia, during early development and/or in slice cultures. However, microglia synapse interaction has not been assessed in the hippocampus of adult awake mice. In contrast to previous results, we observed higher contact rates of microglia with dendritic spines than described for cortical areas or slice cultures. We found that average contact rates of microglia with stable spines were increased in awake compared to anesthetized mice. Smaller microglia contact rates of dendritic spines shown by previous studies might be due to the usage of anesthesia, or the nature of *in vitro* approaches.

### Microglia contact rate associated with stability of dendritic spines

Structural plasticity of dendritic spines involves two main events, the loss of existing spines and the emergence of new spines called spinogenesis. We investigated both, spine loss and gain in relation to previous microglia contacts. Microglia are important players during early CNS development especially for removal of supernumerary synapses^16, 17, 45^, but their role for synapse loss in the adult brain is not well understood. Our time-lapse imaging revealed an association between microglia contacts and loss of dendritic spines. If however microglia actively phagocytose dendritic spines, as suggested by previous studies^10, 32, 45^ or rather induce a retraction into the dendrite remains unclear. Weinhard *et al*. recently showed in an *in vitro* model of early development that microglia do not phagocytose postsynaptic synapses but refine presynaptic boutons and axons^14^.

Furthermore, we analyzed the formation of new spines with respect to microglia interactions of the dendritic element, where new spines emerged. Similar to the results for spine loss, we found an association between dendritic spine formation and dendritic contacts by microglia. This finding is in line with previous studies showing the importance of microglia for synaptogenesis^11, 12, 14, 46^. As microglia were shown to display a high degree of brain region-dependent diversity^15^, there is an unmet demand for studies on functional properties of microglia especially in the hippocampus. Here we provide evidence for a link between microglia-dendrite interactions and new spine formation in the adult hippocampus.

Mechanisms for microglia-mediated spine loss presumably involve phagocytosis or trogocytosis as discussed above. However, during our 45min time-lapse imaging we did not detect a single spine loss or spine gain, strongly suggesting two separate timelines, one for microglia motility/contacts and one for spine gain/loss. While microglia contacts and motility take place on the second to minute timescale^10, 49^, structural changes of dendritic spines happen within days^21, 34, 36, 37^. The mechanisms that reconcile microglia contact rate with structural plasticity of dendritic spines remain to be discovered.

### Clustering of dendritic spines and microglia contact rates

By organizing synapses in clusters their activation can generate relatively large action potentials that do not correlate with the total number of single synapses^50, 51^, described by a nonlinear model of signal transduction^52, 53^. This fascilitates information transfer and increases efficiency of signal propagation. A common theory about these clusters is that memory is stored in small and overlapping distributed populations of neurons, in which synaptic clusters on dendritic branches code for conjoint information about time, space and context^52, 54–57^. Previous studies could show preferential turnover of dendritic spines in spatial clusters along dendrites of excitatory cells in the somatosensory, motor, visual and retrosplenial cortex^29–31^. We observed that approximately 55% of dendritic spine turnover was spatially clustered. Microglia contacted spines that emerged or disappeared in clusters more frequently than single gained or lost spines. This might indicate an involvement of microglia in clustered formation and elimination of dendritic spines in the hippocampus.

In summary, this study supports the hypothesis of microglia induced spine formation and elimination in the hippocampus of adult mice. Furthermore, our data demonstrate a link between microglia motility and neuronal activity, which has implications for microglia studies in *in vivo* systems concerning the utilization of narcotics. Moreover, we propose a link between microglia-spine interaction and structural plasticity of dendritic spines in the hippocampus of adult mice. A deeper and profound understanding about the function and the underlying mechanisms of microglia-synapse interactions bears the potential to discover novel therapeutic strategies for diseases in which these are crucial.

## Methods

### Animals

Mice were housed in an animal facility of the German Center for Neurodegenerative Diseases. They were group housed and separated by gender with a day/night-cycle of 12h. Water and food was accessible *ad libidum*. CX3CR1-GFP knock-in mice carry the international name: B6.129P-CX3CR1^tm1Litt^/J (Stock: 5582) Jackson Laboratory. The mouse line expresses „green fluorescent protein" (GFP) in monocytes, dendritic cells, NK cells and microglia cells in the brain. The *Gfp* gene is introduced as a knock-in into the allele of the *Cx3cr1* gene (chemokine C-X3-C motif receptor 1). These mice were first described by Jung *et al*.^19^. YFP-H mice (B6.Cg-TgN(Thy1-YFP-H)2Jrs, Stock: 3782, Jackson Laboratory) express yellow flourescent protein (YFP), a spectral variant of GFP. The protein is expressed in pyramidal cells of the hippocampus and lamina V, *Stratum pyramiale internum* of the cortex. Moreover YFP-expression is found on the surface of sensory cells in dorsal root ganglia, neurons in retinal ganglia and Mossy fibers in the cerebellum. This mouse line was first described by Feng *et al*.^20^. Littermates of both sexes from a heterozygotes crossing of CX3CR1-GFP and Thy1-YFP-H were used for experiments when being around 7-10 month of age.

### Hippocampal window implantation

To access the dorsal region of the hippocampus mice were unilaterally implanted with a hippocampal window as described before^21^. Surgical tolerance and anesthesia was established with an i.p. injection of ketamine/xylazine (0.13/0.01 mg/g body weight). To prevent oedema and excessive inflammation of brain tissue, dexamethasone was administered (0.2 mg/kg s.c.). For analgesia, buprenorphine (0.05 mg/kg s.c.) was injected shortly before the surgery. After surgery the analgesic was applied three times daily for three consecutive days to prevent post-interventional pain and related stress reactions. Surgeries were performed 4-6 weeks prior to the first imaging session in order to allow the animal to recover from the procedure and to let the reactive gliosis and inflammation subside. This and all procedures described were in accordance with the intern regulations of the DZNE and animal experimental protocols approved by the government of North Rhine Westphalia.

### *In vivo* imaging in anesthetized and awake mice

Mice were anesthetized using 5% isoflurane, which was subsequently reduced to 1% after the righting reflex had subsided. The mouse was fixed to a custom made headholder and mounted under the microscope on a heating plate (37°C) to keep the body temperature at optimum. Images were acquired with a Nikon 16x water immersion objective (NA 0.8) with a working distance of 3 mm. For awake imaging we used a spinning disk made of styrofoam (r = 10 cm, h = 7 cm), which allowed for imaging of running, head fixed mice that control the speed of the spinning disk independently. Time-lapse images were acquired at 100 μm x 100 μm in xy-direction with a pixel size of 0.088 μm/pixel. To allow correction of movement-artifacts oversampling in z was performed. Z-step sizes were between 0.2 and 0.4 μm. Time-lapse images were acquired every 5 minutes for a period of 45 minutes. Image acquisition was carried out on an upright TrimScope II (LaVision Biotech) equipped with a Ti:Sa laser (Cameleon Ultra II, Coherent) that was tuned to 920 nm to allow for simultaneous excitation of GFP and YFP with a maximum output power of 50 mW to prevent photo damage. GFP fluorescence was acquired with a 480/40 BP Filterset. YFP was separated from GFP by a 510 LP filter and detected using a 535/30 BP Filterset.

### TTX-Application

A 50 μM TTX solution in NaCl0.9% was prepared and stored at −20°C. In order to topically apply TTX on the hippocampus, the hippocampal window was removed under anesthesia. Subsequently the TTX solution was administered with a pipette onto the alveus and incubated for 30 minutes. After the incubation a new, sterile hippocampal window was implanted. The anesthetized mouse was head-fixed under the microscope and imaging started after the narcotic had worn off.

### AAV injections

Mice were anesthetized as described before with Ketamine/Xylazine. The bregma was carefully exposed by removing parts of the skin with an incision over the skull midline. Small holes (ø300 μm) were drilled relative to the bregma (positions see below) and the dura was opened. Mice received unilateral stereotactic injections of an AAV2-hsyn-DIO-hM4D(Gi)-mCherry and an AAV9-CamKII-Cre (pENN.AAV.CamKII 0.4.Cre.SV40 was a gift from James M. Wilson (Addgene viral prep # 105558-AAV9)) into the CA1 and CA3 region of the right dorsal hippocampus. Coordinates for CA1 injections relative to bregma: AP −1.90 mm ML +1.50 mm DV −1.10 mm. Injections into CA3: AP −2.00 mm ML +2.50 mm DV −2.08 mm. The needle was left at the sites of injections for 10 minutes in total during which 0.5 μL of virus were administered at a speed of 0.1 μL/minute allowing the virus to diffuse into the tissue. The skin over the skull was reattached by suturing and mice were given one week to recover before surgeries were performed. Analgesic regimen was performed with three times daily application of temgesic (0.05 mg/kg bodyweigt) for three consecutive days.

### Local inhibition of neuronal activity in *stratum radiatum*

For silencing of neuronal activity in *stratum radiatum* of the right hippocampus, mice received unilateral stereotactic injections of an AAV2-hsyn-DIO-hM4D(Gi)-mCherry and a CamKII-Cre into the CA1 and CA3 region of the right dorsal hippocampus. In order to activate hM4D(Gi), mice were injected intraperitoneally with 3 μg/g bodyweight Clozapine N-Oxide^58^ dissolved in 1% Dimethyl Sulfoxide (DMSO) 40 minutes prior to the imaging session. As a control, the same mice received the solvent (1% DMSO) as vehicle injections only.

### Histology

The viral expression in the injected mice was immunohistologycally validated. Mice were transcardially perfused with PBS, pH 7.4. The brains were removed and fixed overnight in 4% PFA. 100 μm of free-floating slices were permeabilized in 0.5% Triton-X100 for 1 hour. Subsequently, slices were incubated with a mCherry antibody (1:10.000, rat serum, ThermoScientific) in a blocking reagent (4% normal goat serum, 0.4% Triton 1% and 4% BSA in PBS) over night at room temperature. After washing the samples three times with PBS, a secondary antibody was administered (Alexa Fluor^®^ 594 goat anti-rat, 1:400) in 5% normal goat serum/BSA and incubated for 2 h at room temperature. During the last 30 minutes of incubation Nissl staining was carried out (NeuroTrace®, 1:200). Afterwards slices were washed three times with PBS, mounted with Dako Mounting Medium and covered with a glass cover-slip.

### Confocal microscopy

For confocal imaging a Zeiss LSM700 microscope was used in combination with either a 20x air objective (NA 0.8) or a 10x air objective (NA 0.3). Nissl fluorescence was acquired with a 435/455 BP filterset and and excitation filter at 405 nm. Alexa 594 was excited at 555nm and detected with a longpass filter (LP 560 nm). The pinhole was set to one airy unit.

### Microglia motility analysis

To separate GFP and YFP linear unmixing was performed with the unmixing tool in ZEN2010 (ZEISS). Subsequently, z-stacks were median filtered. Individual microglia cells were identified by scrolling through the stacks at each time point and then cropped using ImageJ. Z-stacks spanning around 25 μm in depth were extracted from the original z-stack. The oversampling in z-direction allowed removing distorted pictures from the z-stack by manually scrolling through the stack and deleting individual slices. The stacks were registered by applying the ImageJ “StackReg” plugin^59^. Finally, a maximum intensity projection was carried out, resulting in a 2D projection of the cell and its branches. All recorded time points were merged into one stack and subsequently aligned using the StackReg plugin again. The time points were pseudocolored in red and green resulting in a picture in which red areas account for lost microglia branches, green for newly gained branches and yellow pixels resembling stable parts. The turnover rate (TOR) of individual microglia processes was calculated as the number of red and green pixels divided by the sum of all pixels within a determined region of interest. To specifically measure process turnover, we subtracted all pixels representing the cell body.

### Dendritic spine counting, clusters and microglia contacts

Protrusions emerging laterally from a dendrite with a threshold of 0.4 μm in size were counted as spines independent of their individual shape. In each animal 4-6 radial oblique dendrites were analyzed with a length spanning 20-60 μm. Spines were identified by manually scrolling through the z-stack of subsequent time points. If a spine fell below the threshold of 0.4 μm it was counted as lost. Newly formed protrusion larger than 0.4 μm, were counted as new spines. We counted 4455 spines in anesthetized mice and 5365 spines in awake mice. The spine density *in vivo* was determined similarly as described before^21, 60, 61^. The gained and lost fraction (*F*_gained_ and *F*_lost_) of spines was calculated by dividing the number of gained spines (*N*_gained_) and the number of lost spines (*N*_lost_) by the number of present spines (*N*_present_; *F*_gained_ = 100 *N*_gained_/*N*_present_; *F*_lost_ = 100 * *N*_lost_/*N*_present_). The general turnover of dendritic spines was calculated as (N_lost_+N_gained_)/2*(N_stable_+ N_lost_ + N_gained_). Physical appositions of microglia processes and dendritic spines were counted as contact by manually scrolling through the z-stack if fluorescence of both cell types was not more than ≤ 0.4 μm away from each other in the same focal plane and was detectable in ≥ 2 focal planes. If a spine was newly formed, the length of its basis was measured, which subsequently was used to measure microglia contacts at the dendritic shaft at d0. To measure baseline contact rates at the dendritic shaft, each dendrite was subdivided into 1 μm segments. Randomly (using Research Randomizer, Urbaniak, G. C., & Plous, S., 2013), 5 of these segments were tested for microglia interaction.

Previous studies already showed preferential turnover of dendritic spines in spatial clusters along dendrites of excitatory cells in the somatosensory, motor, visual and retrosplenial cortex^29–31^. Taking into consideration that by means of two-photon microscopy the detectable spine densities of apical pyramidal neurons in the hippocampus (~1.1 μm^−1^)^21^ exceeds those found in the cortex (~0.4 μm^−1^)^61–63^ and is obviously still underestimated^37^, we applied a stricter definition of spatial clustering than described before. Chen et al. showed spatial turnover in clusters of 10 μm on dendrites in the neocortex^29^. Modified from this study we defined clusters as events of synaptic turnover happening within a maximum range of 4 μm to each other, accounting for the higher spine densities in the hippocampus. We defined three different classes of clustered events: lost, balanced and gained (Supplementary Fig. 3a); If two or more spines were either gained or lost in proximity of less than 4 μm, they were classified as a “gained” or “lost” clustered event. If one spine was lost and another one gained within the 4 μm distance, the event was classified “balanced”.

### Statistics

Quantifications, statistical analysis and graph preparation were carried out using GraphPad Prism 5 and 7 (GraphPad Software Inc., La Jolla, USA). Mice were assigned to experimental groups in an age and sex balanced manner. To test for normal distribution of data, D’Agostione&Pearson omnibus normality test was used for sample sizes of n > 6 and the Shapiro-Wilk normality test for n < 6. Statistical significance for groups of two normally distributed data sets paired or unpaired two-tailed Student’s t-tests were applied. If no normal distribution was evident, Mann-Whitney Test for groups of two was used. One-way ANOVA with Tukey’s or Bonferroni’s multiple comparison test were performed on data sets larger then two, if normally distributed. For comparison of more than two not normally distributed data sets the Kruskal-Wallis test was performed with Dunn’s correction for multiple comparisons. If not indicated differently, data are represented as mean ±SEM. Figures were prepared with Illustrator CS5 Version 15.0.1 (Adobe).

## Acknowledgements (optional)

This work was supported by the DZNE, grants from the Deutsche Forschungsgemeinschaft (SFB 1089 C01, B06), Centres of excellence in Neurodegeneration (CoEN3018) and ERA-NET grants MicroSynDep, Microshiz. We thank P. Thevenaz and Erik Meijering for the development of the ImageJ plugins “stackreg” and “TurboReg”. We acknowledge Bryan Roth for depositing the Addgene plasmid #44362. pENN.AAV.CamKII 0.4.Cre.SV40 was a gift from James M. Wilson (Addgene viral prep # 105558-AAV9). We thank the animal and light microscopy facility of the DZNE for constant support.

## Author contributions

F.N. conducted the experiments, analyzed the data, prepared figures and wrote the manuscript. S.P., L.C.S., M.M., J.S. and K.K. provided technical assistance. M.F. wrote the manuscript, coordinated research and supervised the project.

## Material & Correspondence

Materials are available on request. All correspondence should be directed to Martin Fuhrmann.

## Supplementary information

**Figure S1.**
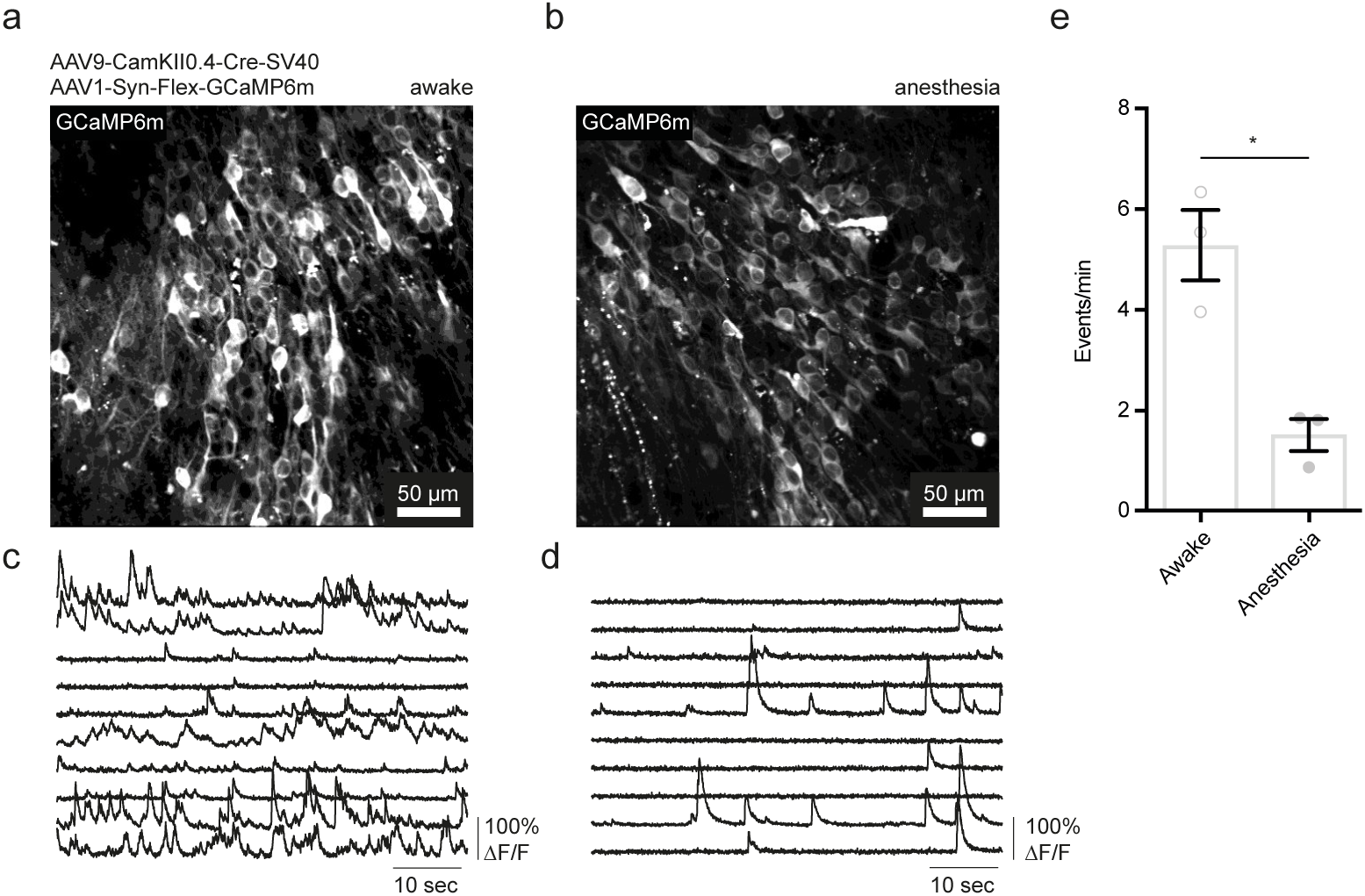
Isoflurane anesthesia decreases neuronal activity in the hippocampus. **a,b** Exemplary image of GCaMP6m expressing neurons in the hippocampus of an awake (a) and an anesthetized (b) mouse (average intensity projection of 1800 pictures acquired within 60 seconds at 30Hz sampling rate). **c, d** Exemplary Ca^2+^-transients from 10 CA1 neurons in an awake (c) and an anesthetized (d) mouse. **e** Average Ca^2+^-transient frequency (events per minute) comparing awake and anesthetized mice. n = 3 mice per group and 50-100 cells/mouse/recording in two different fields of view respectively. Paired t-test, p = 0.0435; * p < 0.05.

**Figure S2.**
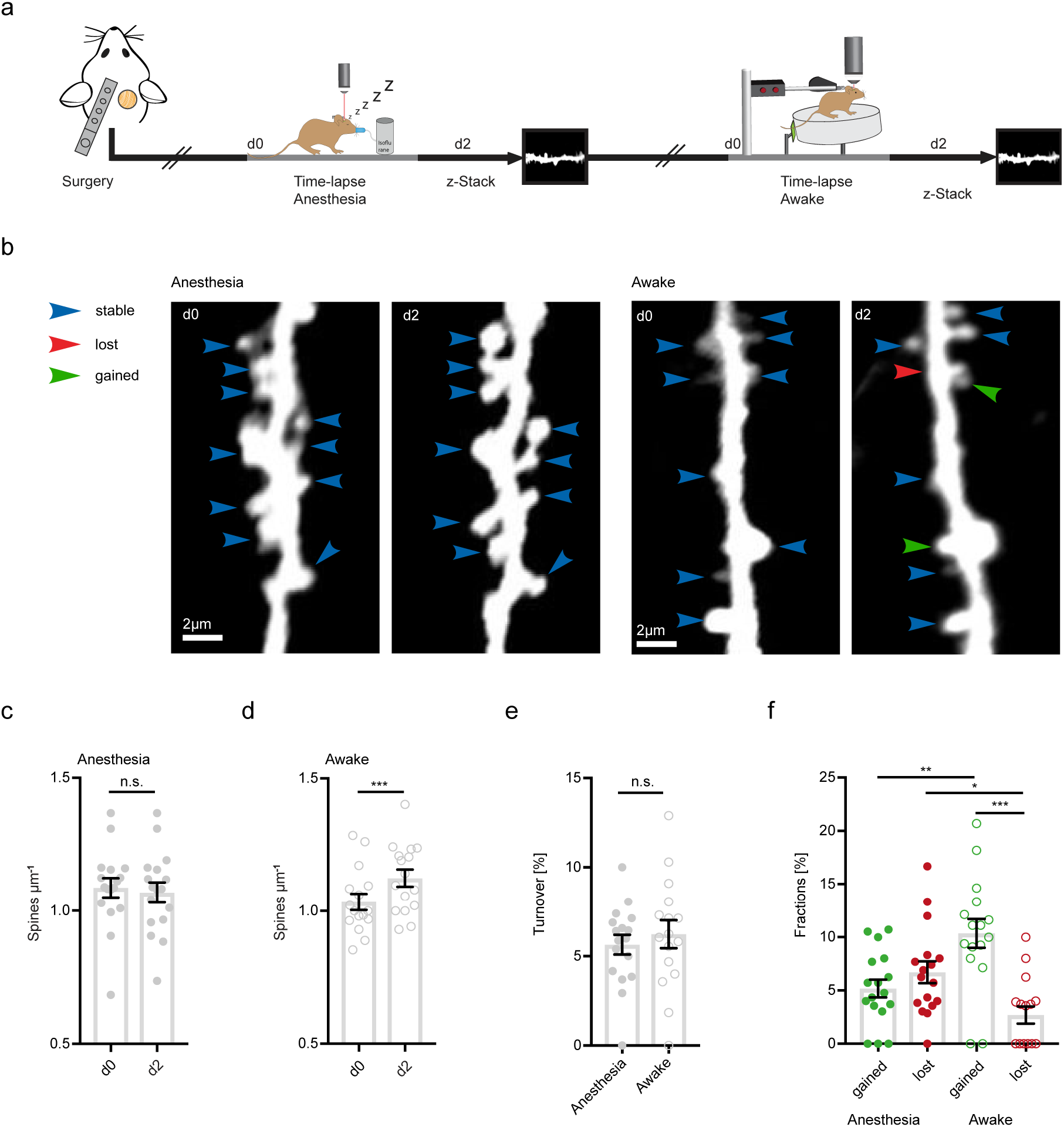
Increased structural plasticity of dendritic spines in the hippocampus of awake mice. **a** Mice were imaged twice with a 2-day interval, first in anesthesia and subsequently awake on a spinning disk. **b** We acquired z-stacks containing radial oblique dendrites in *stratum radiatum* of the hippocampus. Exemplary images of two dendrites acquired with a 2-day interval in anesthesia (left panel) or awake (right panel). Blue arrows indicate stable, red arrows lost and green arrows gained spines. **c** No changes of spine density after imaging in anesthesia (1.09 ± 0.04 μm^-1^ at d0 vs. 1.07 ± 0.04 μm^-1^ at d2; Wilcoxon matched-pairs signed rank test, p = 0.2759). **d** Elevated spine density after awake imaging (1.04 ± 0.03 μm^−1^ to 1.12 ± 0.03 μm^−1^; paired t-test, p < 0.0001). e Similar turnover of dendritic spines was detectable between anesthetized and awake mice (5.64 ± 0.55% anesthesia vs. 6. 23 ± 0.79% awake; unpaired t-test, p = 0.5377). **f** Turnover of dendritic spines differentiated into fractions of lost and gained spines under both conditions. In awake mice the fraction of gained spines was elevated (10.35 ± 1.35% in awake mice and 5.18 ± 0.84% in anesthetized mice; p = 0.0036). These fractions were comparable under anesthetized conditions (gained: 5.18 ± 0.84% vs. lost: 6.7 ± 1.02%; p = 0.7079). The fraction of lost spines was decreased in awake compared to anesthetized mice (Fig, 3f; 2.7 ± 0.81% awake vs. 6.7 ± 1.02% anesthesia; p = 0.0359), ordinary one-way ANOVA followed by Tukey’s multiple comparisons test, F(3,62) = 9,58. Data from 4 mice with spine turnover averaged over dendrites; n = 17 loss + 17 gain in anesthesia and n = 16 loss + 16 gain in awake awake mice respectively (c-f).; n.s. not significant, * p < 0.05, ** p < 0.01, *** p < 0.001, **** p < 0.0001.

**Figure S3.**
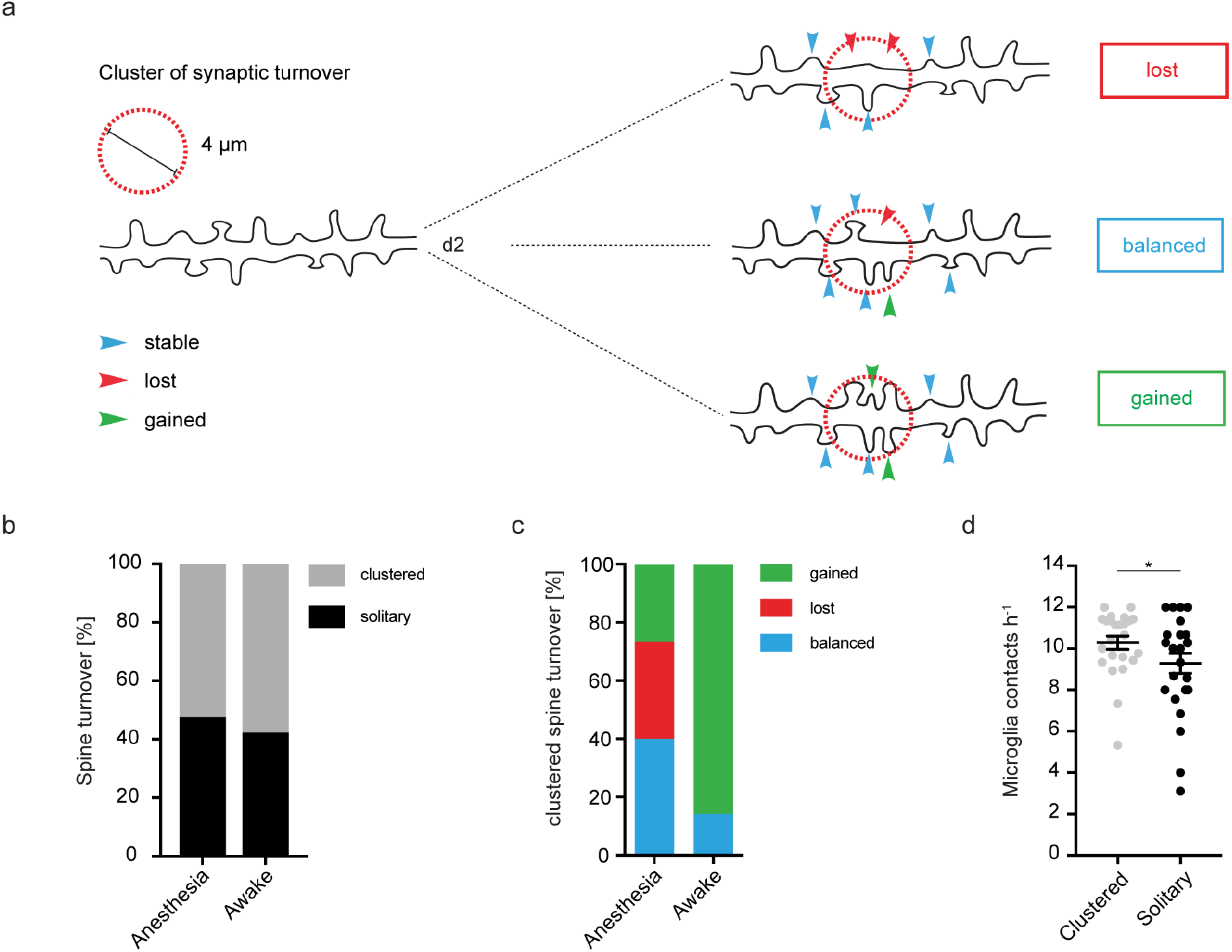
Clustered structural plasticity of spines associated with microglia contact frequency. **a** Schematic, illustrating clustered spine turnover quantification. Three different classes of clustered events (within a distance of 4 μm) were defined. Lost: If two neighboring spines were lost; Balanced: if one spine is lost and another one gained; Gained: if two spines were gained. **b** Approximately 55% of all observed events of structural plasticity were clustered both in anesthesia (52.26%) and awake (57.74%) conditions. **c** Analysis of clustered events with respect to their class (see (a): lost, balanced, gained). Note that in awake mice the fraction of clustered gained spines (85.71%) is larger than in anesthetized mice (26.67%). **d** Microglia contact rates as calculated in Fig.3a preceding a clustered or a solitary event (unpaired t-test, p = 0.0365; * p < 0.05).

**Figure S4.**
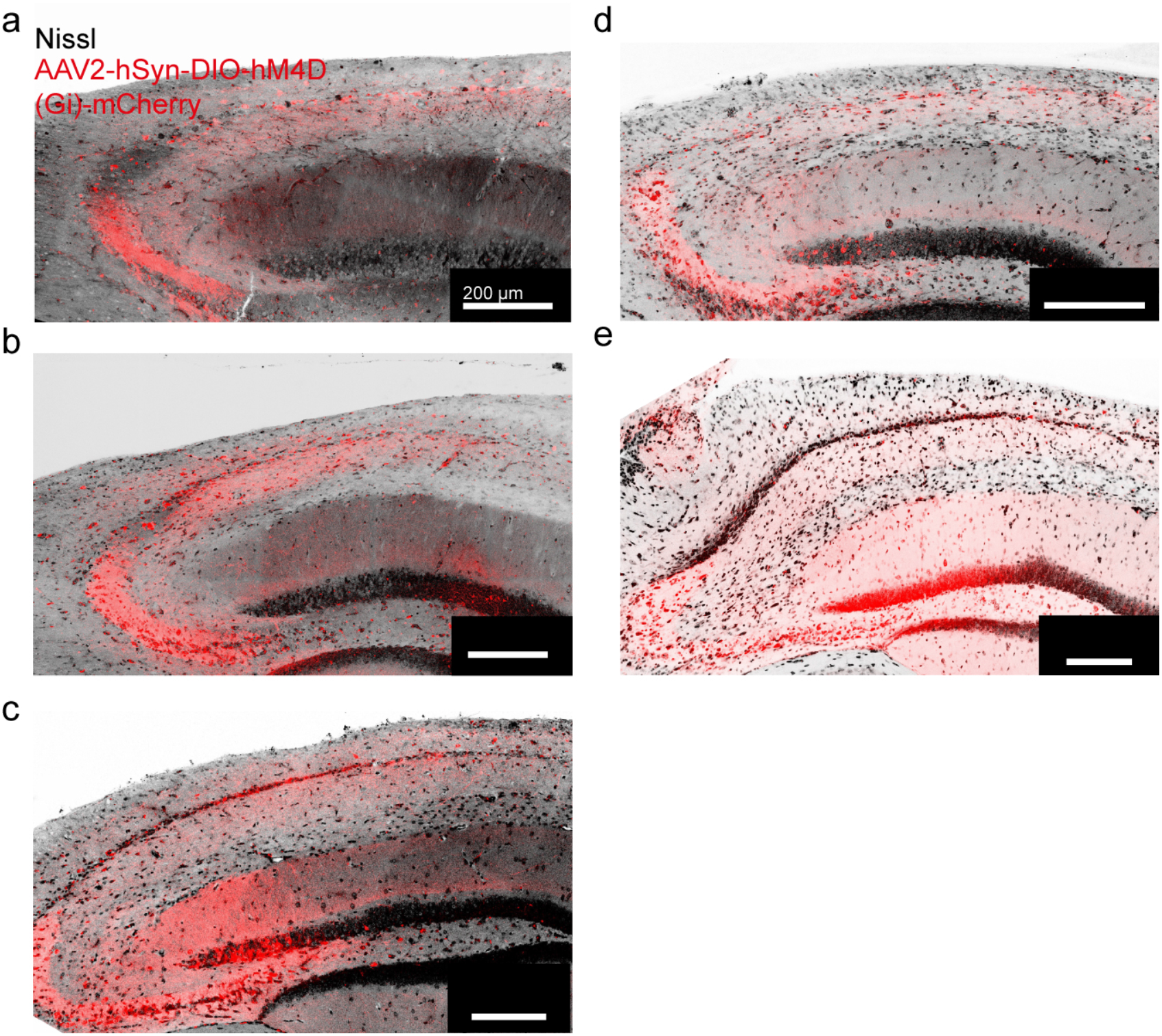
Exemplary expression pattern of DREADDs in the hippocampus of experimental mice. **a-e** Coronal sections of each mouse (5 in total; a-e) used in the experiments were stained post hoc for Nissl (black) and mCherry (red) to check the expression pattern of hM4D(Gi) within the hippocampus.

**Video S1 Microglia motility in an anesthetized mouse**

Microglia fine process motility recorded in the hippocampus of a mouse under isoflurane anesthesia (1% in O_2_) using two-photon microscopy over 45 minutes. Z-stacks were acquired 5 minutes apart each. Scale bar 10 μm.

**Video S2 Microglia motility in an awake mouse**

Microglia fine process motility recorded in the hippocampus of an awake mouse head-fixed on a circular treadmill using two-photon microscopy. Z-stacks were taken 5 minutes apart each. Scale bar 10 μm.

**Video S3 Microglia interacting with neuronal tissue in the hippocampus (awake)**

Microglia (green) engage in contact with the dendritic shaft and individual dendritic spines (magenta). Regions of colocalization of microglia and neuronal structures appear white. Z-stacks were taken 5 minutes apart each. Scale bar 5 μm.

**Video S4 Calcium imaging of CA1 pyramidal cells in an awake mouse**

Exemplary video of Ca^2+^-imaging (GCaMP6m) in CamKII+ cells in the hippocampus of an awake mouse over 60 seconds.

**Video S5 Calcium imaging of CA1 pyramidal neurons under isoflurane Anesthesia**

Exemplary video of Ca^2+^-imaging (GCaMP6m) in CamKII+ cells in the hippocampus of an anesthetized mouse (isoflurane 1% in O_2_) over 60 seconds.

